# Activity Dynamics and Signal Representation in Striatal Network Model with Distance-dependent Connectivity

**DOI:** 10.1101/081752

**Authors:** Sebastian Spreizer, Martin Angelhuber, Jyotika Bahuguna, Ad Aertsen, Arvind Kumar

## Abstract

Striatum is predominantly inhibitory and the main input nucleus of the basal ganglia. A functional characterization of its activity dynamics is crucial for understanding the mechanisms underlying phenomenon such as action selection and initiation. Here, we investigated the effects of the spatial connectivity structure on the emergence and maintenance of localized bumps of activity in large-scale striatal networks (~10,000 neurons). We show that in striatal network model in which the distance-dependent connection probability varies in a Gaussian fashion (Gaussian networks), the activity remains asynchronous irregular (AI) and spatially homogeneous, independent of the background input. By contrast, when the distance-dependent connectivity varies according to a Gamma distribution (Gamma networks), with short-range connectivity suppressed, a repertoire of activity dynamics can be observed: While weak background inputs induce spatially homogeneous AI activity, stronger background inputs induce stable, spatially localized activity bumps as in ‘winner-take-all’ (WTA) dynamics. Interestingly, for intermediate background inputs, the networks exhibit spatially localized, but unstable activity bumps (Transition Activity, TA), resembling the experimentally observed neuronal assembly dynamics in the striatum.

Among the three main regimes of network activity (AI, WTA, TA) we found that in the AI and TA regimes, network dynamics are flexible and can be easily modified by external stimuli. Moreover, the dynamical state of the network returns to the baseline after the stimulus is removed. By contrast, the dynamics in the WTA state are rigid and can only be changed by very strong external stimuli. These results support the hypothesis that the flexibility of the striatal network state in response to stimuli is important for its normal function and the ‘rigid’ network states (WTA) correspond to brain disorders such as Parkinson’s disease, where the striatum looses its repertoire of dynamic states and is only receptive to very strong inputs.

## Introduction

The basal ganglia (BG) is a collection of nuclei located at the base of the forebrain and involved in a variety of functions, including voluntary movement control, decision making, procedural learning and a variety of cognitive and emotional functions. Pathological changes in any of the BG nuclei may lead to brain disorders such as Parkinson’s disease (PD), Huntington’s disease, Tourette syndrome, impulsivity and obsessive-compulsive disorders. Striatum is the main input stage of the BG and therefore, plays a major role in BG related brain function and dysfunction.

Like many other sub-cortical structure, the striatum is a network of inhibitory neurons driven by excitatory inputs from all regions of the neocortex [1], thalamus and hippocampus. The output of the striatum directly projects to the globus pallidus (GP) both internal and external segments and form two of the integral functional pathways of BG, i.e. the Go and No-Go pathways, respectively. An interaction of these pathways form the basis of functions such as action-selection, initiation and execution. The mechanistic details of how a sparsely connected network of inhibitory neurons like the striatum performs multi sensory, sensori-motor integration and action-selection remain poorly understood. In the No-Go pathways, striatum needs to alter the activity of the GP which typically spikes at ≈ 40Hz [2].

Experimental data suggests that in the ongoing activity striatal firing rates are very low (≈ 1 – 3 Hz) [3, 4] and in a given task, up to 30% of striatal projection neurons respond with a markedly increased firing rate (≈ 10 – 20 Hz) [4, 5]. However, it remains unclear, how the task-related activity of the striatum is organized in space and time to shape the activity of GP to initiate action-selection.

Recently, the observation of temporal clustering of spiking activity in the striatum in a decision making task [6] suggest that the network activity may be organized in transient co-activations of small neuron groups, often referred to as neuronal assemblies (NA). Recent *in vivo* imaging studies have shown that NAs are spatially localized and can encode locomotion relevant information in the dorsal striatum [8]. These observations corroborate similar findings on the organization of striatal activity *in vitro* [7]. Co-activated neurons as a part of NAs can effectively increase the influence of striatal neurons on the downstream GP network. However, this observation raises a key question: Under what conditions such NAs appear in the activity dynamics of striatum as a sparse connected, inhibitory recurrent network.

Dynamically, NA type activity state may represent the outcome of a so-called ‘winner-less-competition’ (WLC) [9] or a noise driven transition state between the asynchronous-irregular (AI) activity state and the ‘winner-take-all’ (WTA) state. While the mechanisms underlying the emergence of such transient NA are not well understood, a heterogeneity in both, the external input and in the mutual connection probabilities among striatal projection neurons are common properties of various models that have been proposed to explain the emergence of transient NAs [10–12]. However, these phenomena is usually reported in relatively small networks. It is not clear whether large-scale networks can also exhibit NAs.

In this work we investigate the existence of NAs in a large-scale network model of striatum. Deviating from the previous work that used a small randomly connected networks, we focus on a large-scale network in which neurons are connected in a distance dependent manner. To better understand computations and the encoding and processing of information in spatially connected inhibitory networks we address two key questions: (1) How does the structure of the network connectivity define the repertoire of the ongoing activity dynamics? (2) Under which conditions can task-related inputs modify these ongoing activity dynamics?

To face these questions we investigated the effect of the spatial connectivity structure on the emergence and maintenance of spatio-temporally clustered activity in large striatal network. Specifically, we considered two different types of spatial connectivity profiles in which the connection probability between any two neurons varied as a function of the distance between the neurons, according to either a non-monotonic Gamma distribution or to a monotonic Gaussian distribution [13]. We show that for a non-monotonically shaped connectivity kernel spatially structured activity in the network can emerge. We found that only networks with a Gamma-distributed shaped connectivity kernel can exhibit at least three different activity dynamics regimes. Weak background inputs and/or high input variance induced unstructured AI, whereas stronger background inputs and/or low input variance induced a stable dynamics (e.g. WTA). In an input regime close to the noise threshold the network activity organized into unstable spatial bumps (transition activity, TA), resembling the experimentally observed NAs [6] and WLC [14]. Moreover, the dynamic state of the bump activity also determined the impact of external stimuli on the network activity dynamics. We show that the TA state the network activity is highly sensitive to external inputs both in terms of changes in the average firing rates and pair-wise correlations. In the TA state, the stimulus resulted in a wide spectrum of positive and negative correlations which would be required to initiate plasticity processes to learn the stimulus structure.

Thus, we propose that a non-monotonically shaped (e.g. Gamma distribution) distance-dependent connectivity offers a rich dynamical activity repertoire in purely inhibitory networks such as the striatum, enabling the ongoing activity to dynamically switch between different network activity states. These results will help us to better understand computations and information processing in striatum and other sub-cortical brain regions which are primarily inhibitory networks. In addition, they may help reveal underlying mechanisms of brain diseases such as Parkinson’s disease, in which the basal ganglia lose their flexibility for encoding cortical input stimuli.

## Results

Neuronal activity in the striatum is maintained by thalamic and cortical excitatory inputs. In a randomly connected inhibitory network model of striatum, the recurrent inhibition and the level of external excitatory inputs define the dynamical states and stimulus response properties of the network [11,12,18,23]. Here, we extend this earlier work by investigating the effect of spatial connectivity on the dynamical states and stimulus-response properties of striatal network models. To this end, we used both neural field equations and numerical simulations of large-scale inhibitory networks with 10,000 spiking neurons.

In our network the profile of the spatial connection probability between any two neurons could vary either monotonically or non-monotonically as a function of the distance between neurons. Here, we used two different kernels for spatial connectivity: In *Gaussian networks* the connection probability decreased monotonically in a Gaussian manner as a function of distance (on-center inhibition [13], Fig. 1a, top), whereas in *Gamma networks* the non-monotonic distance-dependent change in connection probability was modeled as a Gamma distribution (off-center inhibition [13] Fig. 1a, bottom).

**Figure 1.**
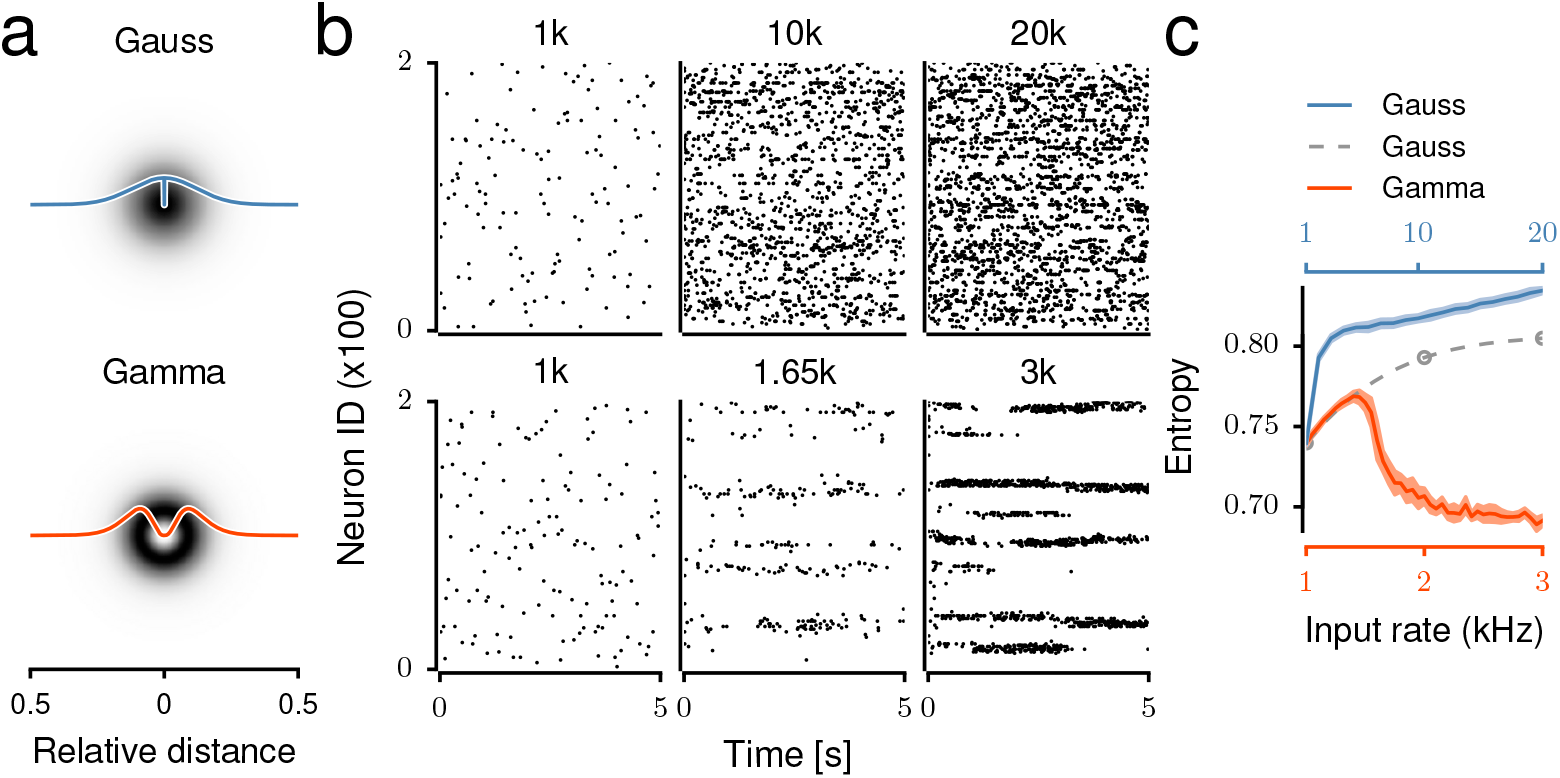
Spike train activity of the network with Gaussian (top) and Gamma-distributed (bottom) connectivity. **(a)** Connection probability between neurons as a function of the distance between two neurons. **(b)** Examples of spiking activity of 100 neurons are visualized for different background input strength. With a higher background input rate the spiking activity of the *Gaussian networks* remained irregularly and the bursting behavior increased. The *Gamma networks*, on the other hand, yielded transient or persistent bursting behavior and local synchronization of spiking activities. **(c)** The informative content of the signal is characterized by the entropy of the neural activity. Different colors of curves refer to the different scales on the abscissa. The gray, dashed curve represents the entropy of the *Gaussian networks* but on input scale of the *Gamma networks* to compare with its entropy (red curve). With an increase of background input rate, the entropy of the population activity in *Gamma networks* showed a drop, indicating an increase in relevant information in this regime.

### Firing rates and spike train irregularity

First, we determined the firing rate and the degree of irregularity of the spiking activities in the networks connected according to the Gaussian (Figs. 1a,b; top) and Gamma (Figs. 1a,b; bottom) shaped connectivity profiles.

As expected, the average firing rates in both network types increased as a function of the background excitatory input (*ν*_*ext*_) (Fig. 1b, top and bottom rows). However, *Gamma networks* were clearly more sensitive to a change in the background input rate and the same average firing rate as in *Gaussian networks* was achieved already for much lower input rates (Fig. 2a, top row).

**Figure 2.**
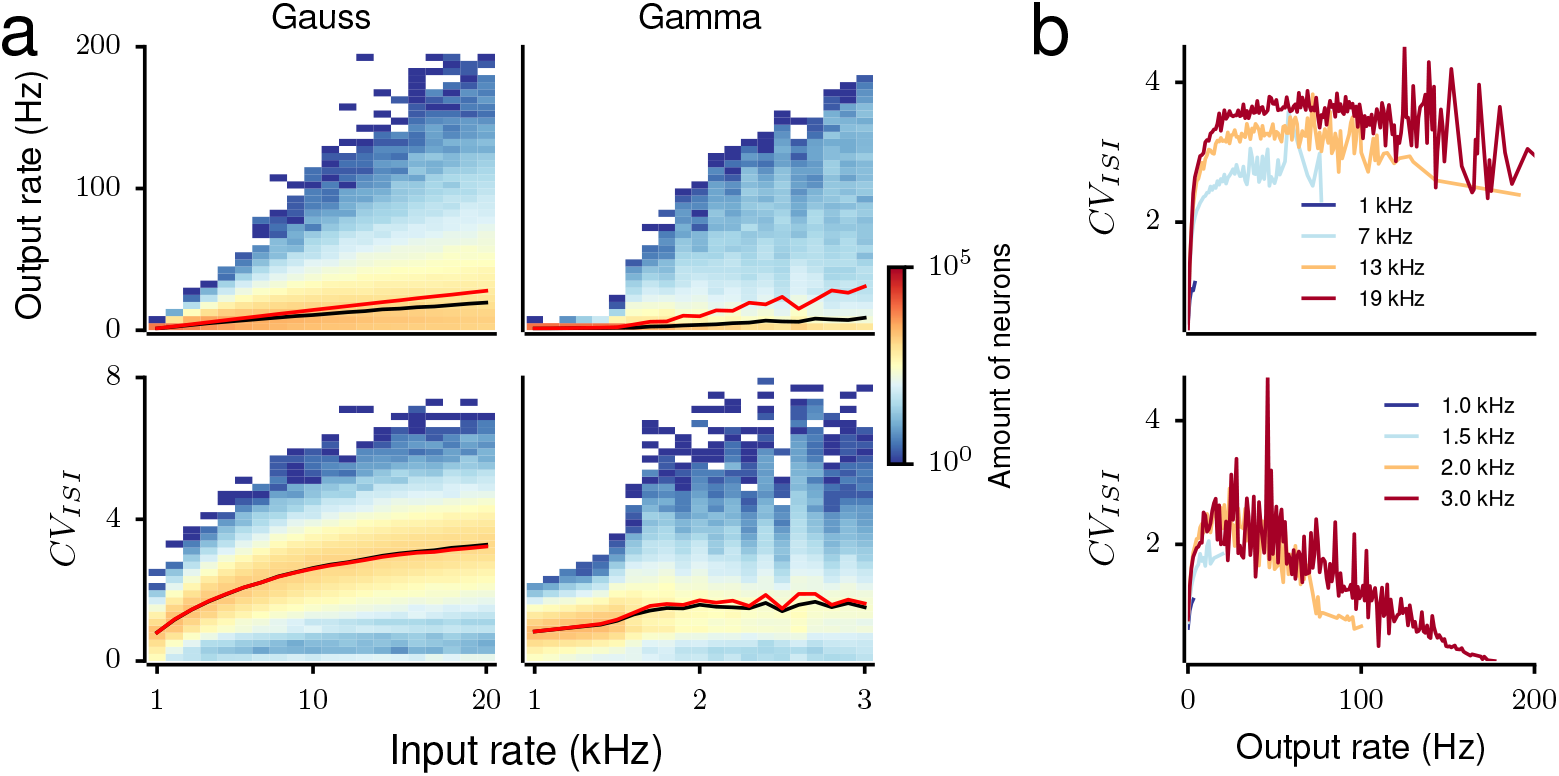
Analysis of the spike patterns in networks with different connectivity profiles. **(a)** Distribution of the mean firing rates (top) and coefficient of variation (bottom) as a function of background input strength for Gaussian (left) and Gamma (right) networks. In *Gaussian networks*, increasing the background input results in a steadily widening distribution of mean firing rates and *CV*_*ISI*_, which for a large fraction of neurons tends to the value 3. By contrast, in *Gamma networks* the distribution of mean firing rates and *CV*_*ISI*_ is rapidly widely distributed from a relatively low input strength (1.5 kHz) upwards. *Gamma networks* are clearly more excitable than *Gaussian networks*. Black curves refer to the median value of the firing rate and *CV*_*ISI*_, respectively; red curves refer to the mean value. **(b)** Relationship between the irregularity of spiking pattern and the mean firing rate. The color of the curves represents the background input rate. In *Gaussian networks* (top) neurons with higher firing rate tend to exhibit a higher *CV*_*ISI*_, whereas in *Gamma networks* (bottom) they tend to exhibit a lower *CV*_*ISI*_.

Moreover, the two network types differed considerably in the distribution of firing rates over neurons (Fig. 2a, top row) and the spike timing irregularity (*CV*_*ISI*_) distributions (Fig. 2a, bottom row). The recurrent inhibition in the network ensured that even for very strong background inputs only few neurons could spike at high rate and a major fraction of neurons spiked at relatively low rates. As a result, increasing the rate of background input to these networks increased the width of the firing rate distribution, this widening being more pronounced in *Gamma networks*.

In *Gaussian networks* the firing rate distribution became distinctly less skewed when the rate of background input was increased, in *Gamma networks* this was hardly the case (Fig. S1). In *Gamma networks*, the firing rate distribution widened drastically at around 1.5 kHz input rate (Fig. 2a top right), thereby increasing the average firing rate. Note, however, that the median output firing rate in *Gamma networks* remained at levels comparable to that in *Gaussian networks* (Fig. 2a, top row).

Interestingly, in *Gaussian networks* the neurons’ *CV*_*ISI*_ monotonically increased with an increase in their firing rate. An average *CV*_*ISI*_ > 1 indicates that most high firing rate neurons spiked in bursts (Fig. 2b,top). Because we used leaky-integrate-and-fire neurons, the bursting in the network activity was caused by the recurrent inhibition and not because of the intrinsic properties of the neurons. In the *Gaussian networks* the connection probability peaked only over a small range in the vicinity of a given neuron, thereby reducing the effective recurrent inhibition and allowing the neuron to maintain its high firing rate for some time. However, fluctuations in the background input (modeled as Poisson spike trains) could rapidly switch the high-rate activity from any one neuron to another, thereby, creating a spike pattern consisting of short-lived bursts and pauses.

By contrast, in the *Gamma networks* only neurons with moderate output firing rates showed a *CV*_*ISI*_ > 1 (Fig. 2b, bottom). Neurons with very high firing rates (*Fr*_*out*_ ≥ 100 Hz) spiked in a regular manner (*CV*_*ISI*_ ≤ 0.5). Such small *CV*_*ISI*_ implies that the network was operating in the so called ‘mean-driven-regime’ [24], in which the background input is strong enough to keep the free-membrane potential of the neurons above spiking threshold. Within the physiological range of output firing rates (≤ 50 Hz), neurons in both network types elicited spike bursts, but the *CV*_*ISI*_ in *Gaussian networks* was nearly twice as high as that in *Gamma networks*. Hence, a Gamma shaped spatial connectivity kernel increased the sensitivity of the network activity to background input and gave rise to a wide range of output firing rates.

### Spatial structure of the network activity

Next, we included the spatial information about neurons into our analysis and characterized the spatial activity patterns in both network types. Visual inspection of the spike rasters (Fig. 1b) suggested that in *Gaussian networks* the structure of the network activity remained spatially homogeneous, even for very high background input rates. By contrast, when increasing the background input rate in *Gamma networks* the activity got confined into local clusters, resulting in spatially periodic activity bumps. To quantify the homogeneity of the spatial activity patterns we measured their entropy (*H*_*S*_) (see Methods). For *Gaussian networks*, the entropy *H*_*S*_ monotonically increased with the background input rate (Fig. 1c, blue trace), indicating that in these networks the activity did not organize into spatial pattens, even for very strong inputs.

By contrast, when the connectivity profile was non-monotonic (here: Gamma distribution) the spatial entropy *H*_*S*_ changed non-monotonically as a function of the background input rate (Fig. 1c, red trace). For a narrow range of weak excitatory input rates (*ν*_*ext*_ ≤ 1.5 Kspikes/sec) the entropy *H*_*S*_ increased from the baseline level observed at the minimum input rate required to obtain any spiking activity in the network. For stronger background inputs (ν_*ext*_ > 1.5 Kspikes/sec) the entropy decreased sharply and reached its minimal value at *ν*_*ext*_ = 3 Kspikes/sec (the highest input rate tested).

This reduction of entropy in *Gamma networks* implies that the network activity became confined to spatially localized regions (Fig. S2, Fig. 3a). Depending on the strength of the background input, three qualitatively different network activity states were observed. In the asynchronous-irregular (AI) state neurons spiked at a low rate and the activity was more or less homogeneously distributed across the network (Fig. S2, top row). For very strong background inputs, the network activity organized into a spatially periodic and temporally stable bump structure (Fig. S2, bottom row). This state resembles the k-winner-take-all (WTA) state [25]. In between, for moderate inputs, the bump structure was aperiodic and unstable (Fig. S2, middle row). We refer to this state as Transition Activity – TA. Recently, Barbera et al. [8] have reported that the activity of D1 and D2 type dopamine receptor expressing striatal neurons is organized as compact spatial clusters. This type of activity of the striatal neurons closely resembles with the TA state.

**Figure 3.**
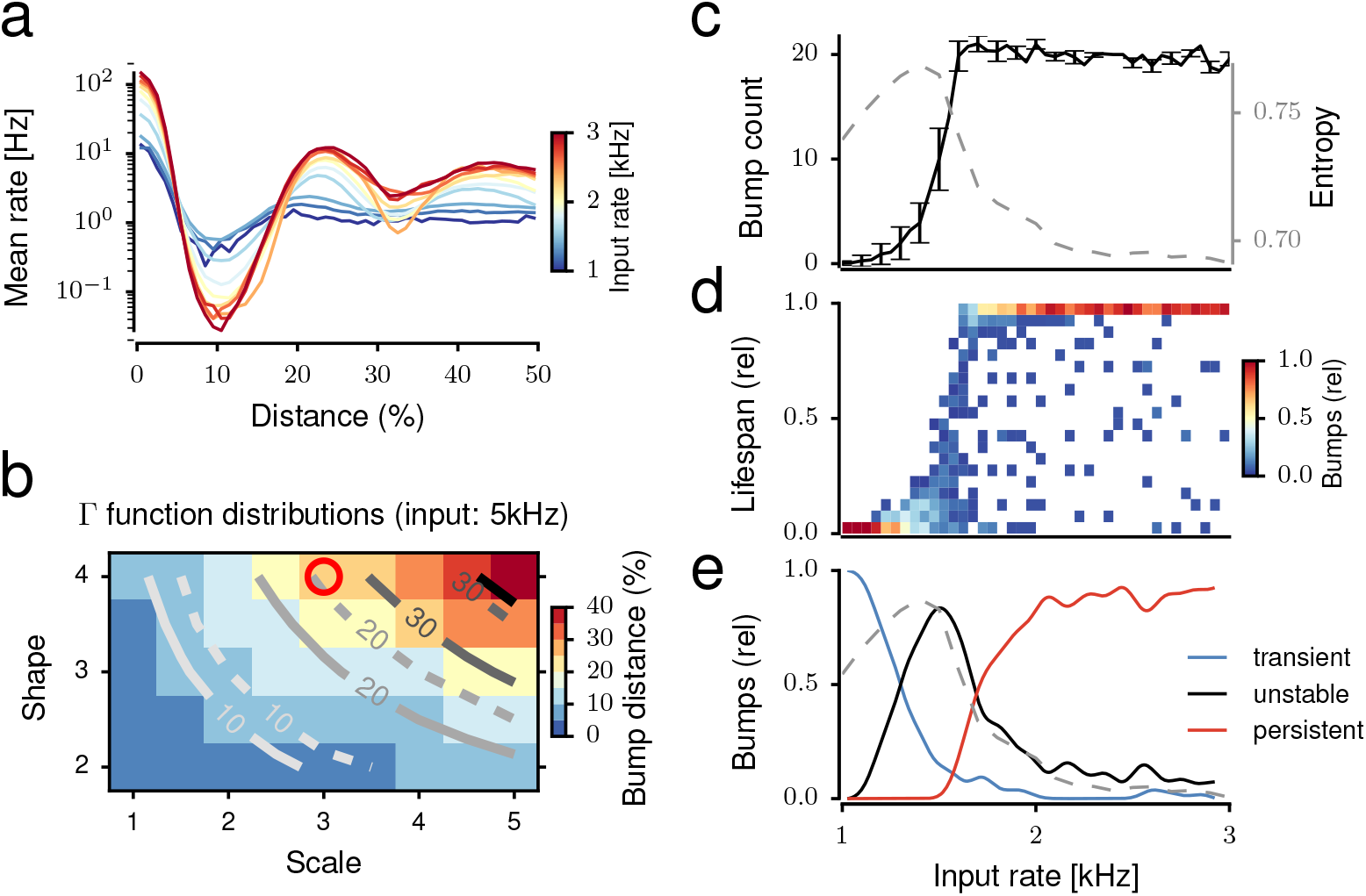
Bump activity in *Gamma networks*. **(a)** Spatial autocorrelation of the spiking activity of the network, showing the mean of the firing rate of each bump depending on the distance from the cluster centroid. Different colors represent the strength of the external excitation. **(b)** Distance between bumps in various Gamma distributions and its comparison in the numerical simulations and when using the analytical methods. The background color is from the network simulation data. Solid curves show results from analytical estimation of bump distance, dashed curves represent the estimation of bump distance from network simulations. The parameter for the Gamma distributed connection is used for the spiking network model (red circle). Spatial representations of activity bumps are also observed in Fig. S3 **(c-e)** The external excitation modulates the nature of the bump activity. **(c)** With increasing background input strength, the number of bumps increases in a sigmoidal fashion. For higher input rates the number of bumps saturates, due to the limited capacity of the finite spatial map. The entropy of the network activity decreases with the increase of the number of bumps and tends to the lowest probability of neuronal activation (dashed gray). **(d)** The lifespan of bumps is related to the dynamic state of the bumps. A shorter lifespan leads to TA dynamics, a larger lifespan leads to stable activity of bumps, creating WTA dynamics. **(e)** The lifespan distribution is split into three groups. The transient (blue) and the persistent (red) appearance of bumps describe AI and WTA states of bump activity, respectively. Between these dynamic states the network is in an unstable state, characterized by a wider distribution of lifespans (black curve). This unstable state of bump activity is related to the entropy of the population activity (dashed gray).

To better characterize the emergence of activity bumps and these three dynamic states in *Gamma networks,* we measured the numbers, sizes, and lifespans of activity bumps. To reliably identify bumps we designed a bump detection algorithm (see Methods). First, we measured the average firing rate as a function of distance from the center of a bump. For weak inputs, the average network activity decayed and reached a baseline level with increasing distance from the center of the bump. However, for stronger inputs not only the firing rate in the bump increased but also the spatially periodic nature of the activity became more apparent (Fig. 3a).

The number of bumps increased in a sigmoidal fashion with the background input rate (Fig. 3c). This increase in bump count was consistent with the decrease in entropy of the spatial activity pattern (Fig. 3c, dashed line). For strong inputs (*ν*_*ext*_ ≥ 1.7 Kspikes/sec), the bumps were stable: neurons that started to spike at the beginning of the simulation remained active for the entire simulation (Figs. 3d and 3e; red trace). For weak inputs (*ν*_*ext*_ ≤ 1.2 Kspikes/sec), the few bumps that appeared lasted only for a short duration (Figs. 3d and 3e; blue trace). For the medium range of input rates (1.2 ≤ *ν_ext_ ≤* 1.7 Kspikes/s), the bumps showed a wide range of lifespans (Figs. 3d and 3e; black trace).

This analysis of the network activity showed that *Gammanetworks* exhibited random unstructured activity for weak background inputs and stable periodic bump activity for strong inputs. The stable bump activity was similar to the periodic bump activity observed in networks of excitatory and inhibitory neurons inter-connected according to a spatial Gaussian connectivity profile [26–28]. This raises the question, why inhibitory networks with Gamma distributed connectivity profile exhibit spatially bump activity pattern and why networks with Gaussian distributed connectivity profile do not?

### Necessary conditions for the emergence of bump states

To address this question we investigated the dynamical states of spatially connected inhibitory neurons using the neural field equations [25,29]. For simplicity we started by formulating the neural field equation for a one-dimensional network with circular boundary conditions. The results hold for a 2-D network with torus-folded boundary conditions because we are considering a homogeneous and isotropic scenario. The mean membrane potential *v*(*x, t*) in the continuum limit is given by:

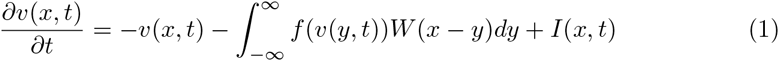

where *W*(*x*) denotes the spatial connectivity profile and *f*(*v*) the neuron transfer function mapping the mean membrane potential to the output firing rate. Without loss of generality we used normalized connectivity profiles *W*(*x*), and the scaling parameter was chosen such that the absolute connection strength was absorbed into *f*(*v*). The background input to the neurons is denoted by *I*(*x,t*). For the network, the stationary and spatially homogeneous solution *v*(*x,t*) = *v*_0_ = *constant* is a solution to the Equation (1) for a constant background input *I*(*x,t*) = *I*_0_*. v*_0_. Here, *v*_0_ is given by the implicit equation:

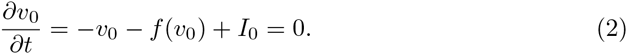

To investigate the stability of this homogeneous solution, we considered small perturbations *v*(*x,t*) = *v*_0_ + *ϵν*(*x,t*) around *v*_0_ and linearized the transfer function in Equation (1). After subtracting Equation (2) this yields

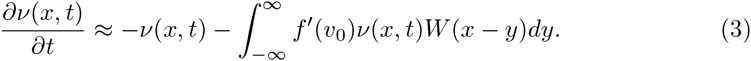

Here and in the remainder *f′* (*v*_0_) is used as shorthand notation for 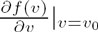. In the Fourier domain, this expression simplifies to:

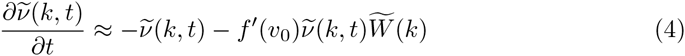

with 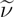 denoting the Fourier transform of *ν* with respect to space. We can now obtain the eigenvalue

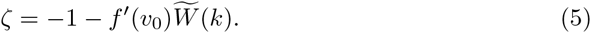

When the eigenvalues are positive, small perturbations do not die out, indicating unstable dynamics. Assuming that the slope of the transfer function (*f′*(*v*_0_)) is always nonnegative, negative values in 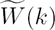 are a necessary condition for positive eigenvalues *ζ* and, hence, for spatially periodic activity bumps. In purely inhibitory networks, this condition can be fulfilled by off-center inhibition kernels, like the Gamma-kernel under investigation, or, e.g., a mixture of two Gaussian distributions arranged symmetrically around zero. While for biologically plausible connection kernels, off-center inhibition, i.e., a non-monotonic kernel, fulfills this condition, it should be noted that this is not a necessary condition. For instance, the Fourier transform of a box-shaped kernel around zero takes negative values at non-zero frequencies and could, therefore, in principle, generate bump states.

For the Gamma-distribution shaped connectivity kernel:

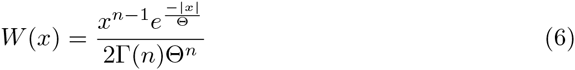

with shape parameter *n* and scale parameter Θ this condition of positive eigenvalue *ζ* > 0 can be fulfilled for *n* > 1, when the Fourier transform 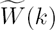 takes negative values (Fig. 4 right panel). By contrast, the Gaussian kernel Fourier-transforms into another Gaussian which never takes negative values (Fig. 4; right panel) and therefore, purely inhibitory networks with a Gaussian type connectivity profile do not show any spatially periodic bump activity. When the eigenvalues *ζ* are negative, the network activity remains spatially homogeneous, similar to the AI state observed in both the *Gaussian networks* and the *Gamma networks* (Fig. 1c).

**Figure 4.**
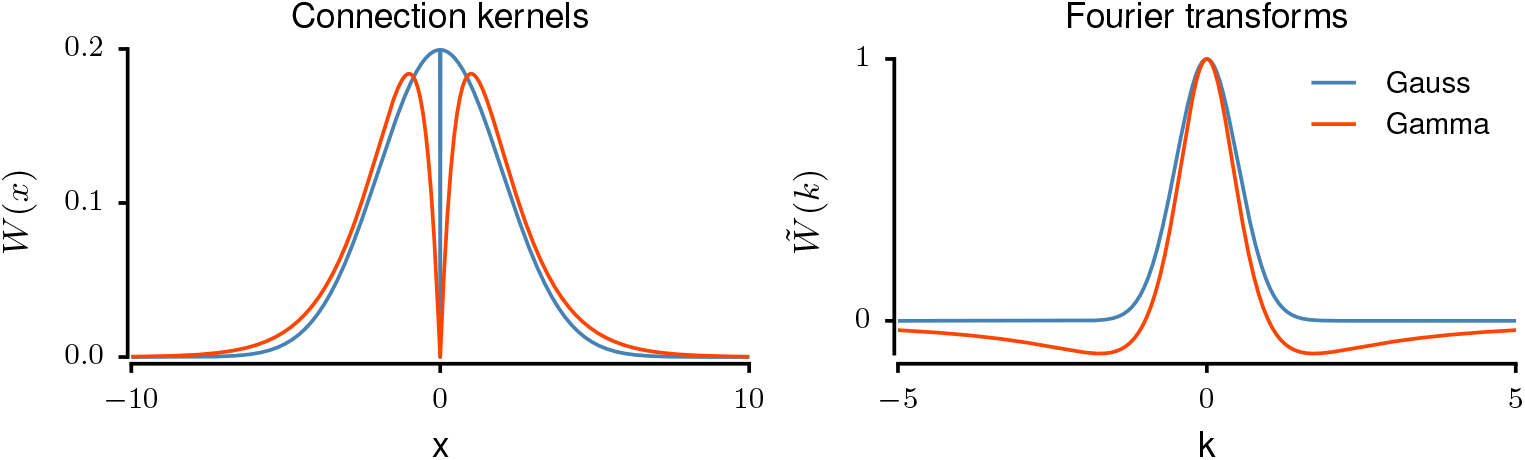
Analytical description of connectivity profiles. The graphs show the spatial connectivity profile (left) and its Fourier transform (right) as a function of the distance between neurons and wave numbers, respectively, for both Gamma and Gaussian connectivity kernels. Note that the spatial connectivity profile remains positive for both connectivity kernels. However, their Fourier transforms behave differently; the Gaussian kernel remains positive whereas the Gamma kernel takes negative values.

Equation (5) also revealed that the slope of the transfer function at stationary rate (*f*′(*v*_0_)) is an important factor controlling the eigenvalues. For positive eigenvalues it needs to be sufficiently large. That is, the following condition needs to be fulfilled:

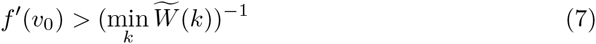

Here, *f*′(*v*_0_) is controlled by a number of factors. For instance, an increase in the synaptic weights increases *f*′(*v*_0_) and can make the network cross the bifurcation. Similarly, increasing the background input will also increase the slope *f*′(*v*_0_), because the transfer function of leaky-integrate-and-fire neurons in the simulated network is convex for low firing rates [30]. That is, both, an increase in the strength of recurrent synapses and an increase in background input rate can cause the transition from spatially homogeneous firing to periodically organized bump solutions.

This analysis shows that the activity of an inhibitory network driven by constant input *I*(*x*) has two stable solutions: for weak inputs the stationary state is spatially homogeneous (AI state, Fig. S2; top row) whereas for strong inputs, the stationary state is spatially periodic (WTA state, Fig. S2; bottom row). When the network is driven with noisy inputs or the neurons have unequal numbers of synapses (due to random connectivity), the transition between the two stable solutions is smoothened (TA state, Fig. S2; middle row). This is consistent with the numerical network simulations (Figs. 1, 3) where noise was introduced into the network by the connectivity and the Poisson type spike trains as external input.

In addition to stating the condition for spatially periodic bump solutions to arise, Equation (5) also enables us to estimate the distance between bumps. The wave number of the emerging spatially periodic solution is approximately given by the wavenumber *k*_*c*_ which minimizes the Fourier transform of the Gamma kernel (6). This critical wavenumber *k*_*c*_ is given by:

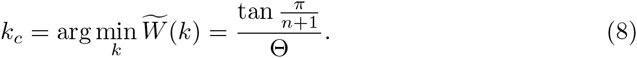

### Properties of the ongoing network activity, required to represent external stimuli

How can the activation and inactivation of bump behavior as observed in the TA or WTA dynamics be useful for information processing in the striatum? Recall that the spiking activity bumps appeared in the absence of any specific stimulus, that is, in response to a non-specific excitation of the entire network. To use a network with such ongoing activity for any processing, it is imperative that the presentation of an external stimulus (i.e. the specific activation of a fraction of neurons in the network) is able to alter the ongoing network state. This requires the ongoing activity dynamics to be ‘flexible’ such that different stimuli can generate different, that is, separable responses – since, evidently, inseparable responses would be quite useless for any information processing. In the ongoing activity observed in TA or WTA dynamics, stimulated neurons could create new bumps and/or attract (or repel) already existing nearby bumps. Subsequently, the stimulus-induced bump could enhance bump formation in nearby neurons by disinhibition. However, the ongoing network dynamics could also be ‘rigid’, meaning that external stimuli would not able to alter the network state. Indeed, in order to participate in a rich repertoire of behaviors the network should exhibit a state in which most neurons can be activated, thereby, allowing the formation of novel spiking bumps with ‘flexible’ dynamics.

Even when the ongoing activity is flexible enough to produce well separated responses to different stimuli, the dynamics could be ‘elastic’, such that after the stimulus is removed the network activity returns to the baseline. Alternatively, it could be ‘plastic’, such that activity does not return to the baseline after the stimulus is removed.

Taken together, we argue that ‘flexible’ and ‘elastic’ ongoing activity dynamics would be optimal for a striatal network to be able to represent and process external stimuli. Therefore, in the next step we investigated the flexibility and elasticity of the ongoing activity dynamics in *Gamma networks* in the AI, TA and WTA states.

### Impact of the ongoing network activity on the stimulus response

To test the flexibility and elasticity of the ongoing network dynamics we stimulated two non-overlapping regions of the network alternately (Stimulus A and B; Fig. 5a, see Methods). In the AI state the network instantaneously responded to the stimulus and switched to a different pattern when the input was changed (Fig. 5a, top row). In this state, there were no bumps in the ongoing activity, therefore, the stimulus created new bumps.

**Figure 5.**
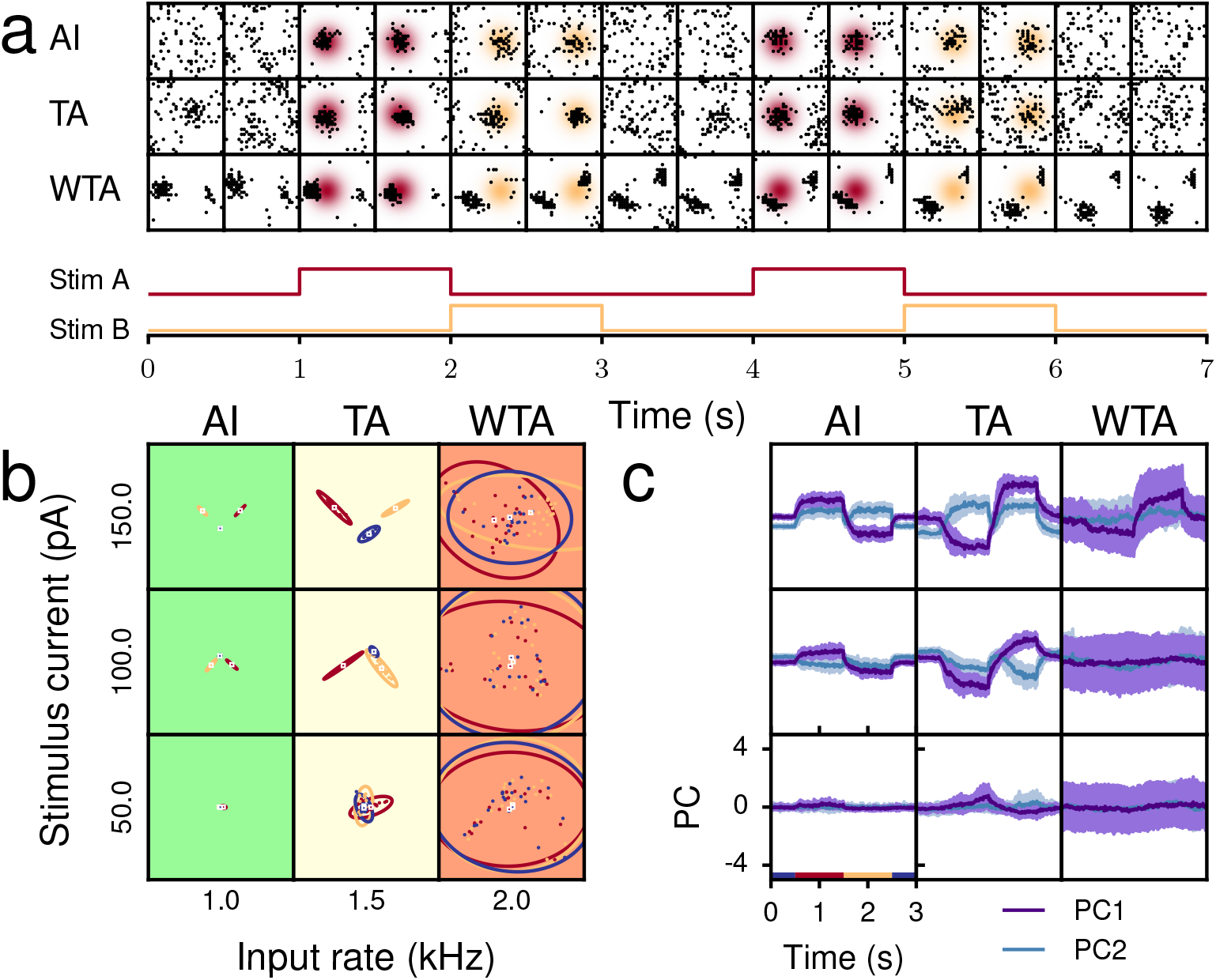
Impact of the network dynamics on the stimulus response with PCA. **(a)** Spatial distribution of spiking activity displayed in time series for different dynamic states (rows). Each block shows a spatial map of 30 × 30 neurons from the ROI (black dots) in a time window of 100 ms, with 500 ms intervals between successive blocks. The gradient background is the probability area for stimulated neurons, its color refers to a stimulus phase. **(b)** Population activity in PCA for different dynamic states of network activity. Each dot represents the principle components describing an activity state of the population in a given time window (1 s). The baseline activity is colored in blue, the other colors represent the activity in corresponding stimulus phases. **(c)** Evolution of the dynamical states over a period (3 s) in different stimulus phases - off-stimulus (blue), stimulus A (red) and B (yellow). Subplots are associated to stimulus currents and input rates in (b). Purple and blue traces represent averaged values of PCs based on time series of the rate histogram with 100 ms bin width over 20 repetitions. Colored areas represent the standard deviation of the PC values over 20 repetitions.

In the TA state, the ongoing state already showed transient bumps. In this state, external stimuli modified the already existing bump pattern to an extent which was different for the stimuli A and B. By contrast, in the WTA state, the ongoing activity showed strong stable bumps and the external stimuli proved insufficient to alter the ongoing bump pattern (Fig. 5a, bottom row).

To test whether the two stimuli elicited unique activity responses that could also be distinguished from the ongoing activity we measured the principle components (PC) of the ongoing and of the stimulus-induced network activity (see Methods; Fig. 5b). The minimum stimulus strength 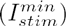 needed to elicit a response discernible from the ongoing activity depended on the ongoing network activity state.

In the AI state, once the input strength exceeded 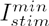, both inputs generated a specific response which clearly differed from the ongoing activity as can be inferred from the well separated PCs of the network activity in ongoing, stimulus A and stimulus B phases (Fig. 5b, columns marked in green).

In the TA state the stimulus threshold 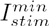 exceeded that in the AI state. However, above this threshold, the network responses to the two stimuli and the ongoing activity again clearly differed from each other (Fig. 5b, column marked in yellow). In the WTA state, however, for all different input strengths we tested we did not observe any response that differed from the ongoing activity (Fig. 5b, columns marked in red). Note that the maximum strength we tested was five times the 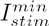 in the AI state. Taken together, these results suggest that among the three regimes of network activity only the AI and TA states were flexible enough to differentially encode different stimuli.

Next, to test for the elasticity of the ongoing activity dynamics we studied the temporal evolution of the first two PCs in the different network states and for different input strengths (Fig. 5c). In the AI state, response switching was nearly instantaneous and after removing either of stimuli A or B the network activity and the two PCs rapidly returned to their baseline values (Fig. 5c; left column). In the TA state, the switching of the network responses was slower, because stimulus encoding in this state involved a reorganization of already existing bump patterns (Fig. 5c, middle column). This could, in principle, be considered a useful property, because such short-term memory of the previous stimulus could be used to encode temporal associations between successive stimuli.

Specific differences in network responses were observed in the temporal correlation over 20 repetitions of stimulus trials (Fig. S4). The spiking activity responses to repeated presentation of the same stimulus were highly correlated in both AI and TA network states, indicating the stability of the network responses (Fig. S4, a,b, II,IV). By contrast, as is necessary for reliably encoding a sequence of different stimuli, the stimulus responses to the two stimuli were uncorrelated (Fig. S4, a,b, III,V). Note that the ongoing activity in the AI and TA state was quite different in different trials and, hence, the temporal correlations in the ongoing activity over trials were very small (Fig. S4, a,b).

In the WTA state, however, the response, if at all different from the ongoing activity, was by far the slowest (Fig. 5c; rightcolumn). Notice also that the variance of the two PCs clearly grew as the activity went from the AI/TA states to the WTA state (Fig. 5b). This increase in variance rendered the separation of responses very difficult. Moreover, the response in the WTA state was not repeatable and temporal correlations between responses to both the same stimulus and to the different stimuli were widely distributed (Fig. S4, c).

Taken together, the stimulus response properties of the three ongoing network activity regimes in *Gamma networks* showed that in the AI and TA states the network dynamics are both flexible and elastic enough to reliably encode different external stimuli arriving in sequence. By contrast, the WTA state proved to be rigid and therefore, not suitable for encoding incoming stimuli.

### Modulation of pairwise correlations is maximal in the TA state

Both AI and TA states are similar in terms of stimulus sensitivity and response separability (Fig. 5). Because, in general, the firing rate in the TA state is higher than that in the AI state and it could be argued that the AI state is a more suitable ongoing activity state for a striatal network. Further analysis of the evoked activity however, revealed a crucial difference between these two states that renders the TA as a more suitable ongoing activity state.

In the AI state, a weak external stimulus only affects the rate of stimulated neurons and thereby induces a very small effect on the neighboring neurons. Therefore, the correlation spectrum during the evoked activity is slightly positively skewed (Fig. 6, left, red trace). That is, while weak stimuli can evoke activity responses in the AI state the spectrum of correlations remains largely unaffected (Fig. 6).

**Figure 6.**
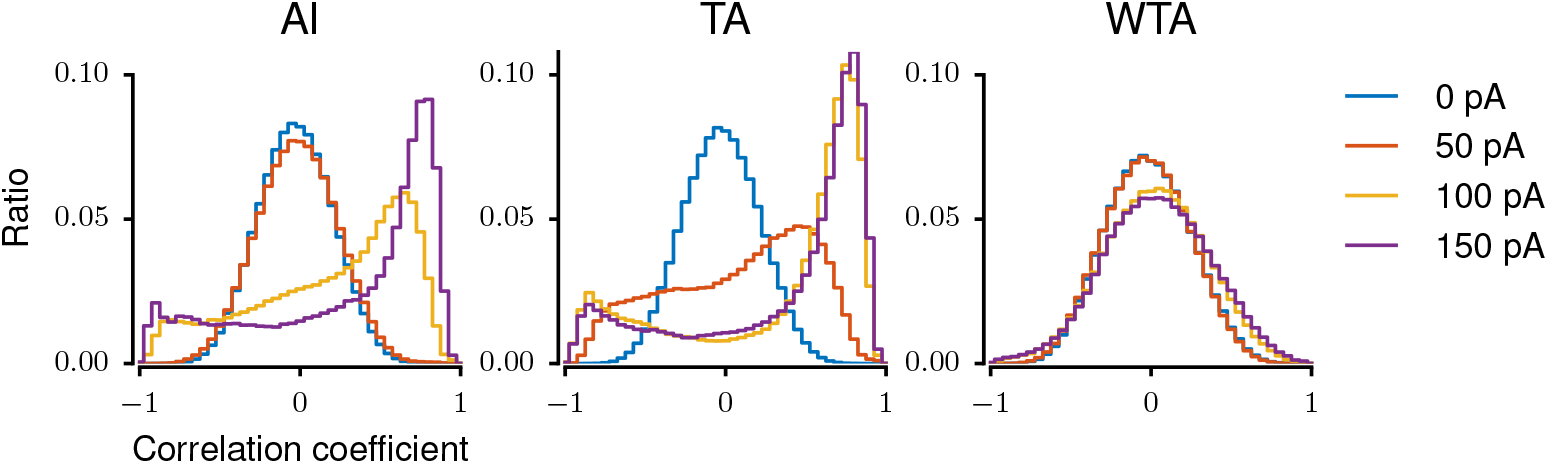
Impact of the network dynamics on the modulation of the correlation spectrum. It displays the correlation spectrum of the activity in three dynamic states (AI, TA, WTA). Colored traces represent the stimulus strength on selected neurons. Compared to correlation spectrum in ongoing activity (blue trace) a higher excitation is required to modulate correlation in AI state than in TA state. With a stronger input the correlation is widely distributed observed in network activity in both AI and TA states. Interestingly, the distribution is not symmetric and the network activity showed more positive correlation than negative correlation.

By contrast, in the TA state, even a weak external stimulus can create new activity bumps and therefore, introduces both positive and negative correlations. (Fig. 6, middle, red trace). The resulting distribution of correlations in the evoked activity is much wider than observed in both AI and WTA states. That is, in the TA state even weak stimuli are able to induce large changes in the structure of pairwise correlations (Fig. 6 middle, red trace). These correlations maybe effective in ‘carving’ out the selected action not only in the striatum but also downstream in GPe and GPi, both of which have high baseline activity and requries coordinated striatal inhibition to be suppressed. For strong external stimulus the correlations spectra are similar in both AI and TA states. In the WTA state a weak stimulus did not induce any visible modulation in the correlation spectrum Fig. 6, right) and only very strong inputs could triggers a small change in the correlation structure (Fig. 6, right, yellow and magenta traces).

Several previous experimental [6,7] and computational studies [10–12] have used the spectrum of pairwise correlations to identify neuronal assemblies in the striatal activity. Our work suggests that the TA state of the striatal network model is more suitable to generate stimulus evoked NAs similar to the ones described previously [6, 7]. Presence of both positive and negative pairwise correlations in the evoked activity implies that different spike timing dependent plasticity (STDP) mechanisms could be exploited to learn stimulus specific NAs. Within the striatum it is still debated whether MSN-MSN synapses exhibit Hebbian or anti-Hebbian STDP [31]. In TA state both positively and negatively correlated pairs exist and therefore, both Hebbian or anti-Hebbian STDP could be used to learn the stimulus-specific NAs. Moreover, both positive and negative correlations allow for a better signal-noise ratio within the striatum as well as passing this signal downstream in the basal ganglia.

## Discussion

Here, we investigated the activity dynamics and stimulus response properties of striatum as purely inhibitory networks with different spatial connectivity profiles. We showed that a non-monotonically changing spatial connectivity profile can lead to the emergence of spatially structured activity in purely inhibitory networks. By contrast, when the connectivity changes monotonically as a function of distance between neurons, the network activity is uniformly distributed over the network irrespective of background input. Specifically, we have shown that with a non-monotonically shaped connectivity profile a striatum network can exhibit three qualitatively different activity states: ‘asynchronous-irregular’ (AI), ‘transition activity’ (TA), and ‘winner-take-all’ (WTA) dynamics. Importantly, among these three different dynamical network states, both AI and TA states have the necessary properties for reliably encoding external stimuli. Finally, we showed that the TA state is an adequate state in which the striatum can support stimulus specific NA and provides the necessary correlation structure to learn such NAs.

Transient neuronal assemblies (NAs) in striatum like purely inhibitory networks were recently defined as groups of neurons showing a conspicuous correlation in their temporal firing rate profiles [10, 11]. Such NAs and their member neurons were identified by an off-line analysis of the spiking activities of neurons in sparsely connected, random recurrent inhibitory network models with weak synapses. In such networks, NAs were found to be randomly distributed over the entire network and appeared to involve mutually unconnected neurons [12]. However, what feature of basal ganglia performs such an offline clustering of spiking activity into assemblies downstream is currently unknown. In the networks we studied, in particular in the *Gamma networks*, potential NAs could consist of groups of neurons, which are unconnected but spatially clustered together and share most of their sources and targets. In the transient TA regime, this would provide a convenient apparatus to rapidly switch their allegiance from one stimulus to the next.

### Relevance for striatal activity dynamics

In its ongoing activity state striatal neurons are relatively silent. Task related activity can increase the firing rates up to 20 Hz [4]. Recently, transient neuronal assemblies (NAs) were reported in striatal projection neurons *in vitro* [7] and *in vivo* [6]. That is, striatal neurons were observed to simultaneously become co-activated and stop being co-activated on time scales in the order of ≈ 100 ms. Moreover, Barbera et al. [8] have also shown compact spatial clustering of the activity of both D1 and D2 type dopamine receptor expressing striatal neurons. Transient NAs were also reported to be modulated by behavioral state, task phase [6, 8] and dopamine level [7]. Beyond the NAs, in a low-dopamine state (as in Parkinson’s disease), striatal neuronal activity lost its diversity and task related switching in neuronal activity was reduced [32, 33].

The three network states that we have identified in this work capture different aspects of the ongoing and evoked activity of the striatum in the normal and low-dopamine states. The low firing rate of striatal neurons and the sparse sampling of their activity in current experiments do not allow us to determine the spatial structure of the striatal network activity in the ongoing state. Both the AI and TA states in our network models match some properties of the ongoing activity measured *in vivo*. For instance, the AI state of the *Gamma networks* closely resembles the ongoing activity state in the healthy striatum, because in this state neurons spike irregularly at low firing rates. The irregularity (*CV*_*ISI*_) is defined by the Poissonian nature of the activity in this state (Fig. 2). Moreover, in this state the network activity dynamics is flexible (Fig. 5) to encode stimuli, which is optimal for its ability to encode.

The transient NAs observed in the TA state in our *Gamma network* models closely resemble the stimulus evoked NAs in the striatum *in vivo* [6]. Further support for this comes from the observation that in behaving rodents neighboring neurons have similar stimulus response properties [4]. Here we obtained TA state by increasing the background input to all neurons in the network. This was done to expose the repertoire of all possible states of the network activity dynamics. It is possible that task-related input from the cortex perturbs the AI state of the ongoing activity and drives the network into the TA state.

We do not want to exclude here the possibility that the TA state could also be the ongoing state of the striatum under some behavioral conditions. TA states offers several useful properties that make it a convenient ongoing activity state to encode stimulus related information. For instance, the transient patterns of NAs offer greater diversity in the ongoing activity for encoding and processing of stimulus-induced information. Furthermore, in the TA state an external stimulus needs to reorganize the existing NA pattern and, likewise, when the stimulus is removed the network slowly relaxes to the ongoing state (Fig. 5c). That is, in the TA state the stimulus response is slower than in the AI state. However, this feature could be used to create associations between successive stimuli or action sequences.

In the WTA state, neurons have higher firing rates and the spatial bumps are stable (Fig. 3; Fig. S2). In this state, the ongoing activity shows only a very small diversity and in this ‘rigid’ state only very strong inputs can induce any response. With these properties, the WTA state closely resembles the striatum dynamics in Parkinson’s disease [32,33]. Moreover, the WTA state is observed when neurons are spiking at a high rate (Fig. 3), which could be achieved either by increasing the ongoing, external excitation or by increasing the excitability of the neurons or reduced lateral inhibition among the striatal neurons. This is also consistent with the fact that in Parkinson’s disease the cortical drive to the striatum is increased due to the potentiation of cortico-striatal synapses [34, 35] and the lateral inhibition among the MSNs decreases [16].

Despite these similarities and interesting insights regarding the striatal activity in the healthy and Parkinson’s disease states, we note that our model is highly simplified and ignores several key features of the striatal network. It would be necessary to study how other components of the striatal network (e.g. fast spiking neurons, separation of striatal network into D1 and D2 type medium spiny neuron populations [36]) affect the flexibility of the stimulus responses in the AI and TA states.

### TA state is optimal for stimulus representation and learning

Going by the criterion of stimulus representation and response separability, both AI and TA states are similar. Because the ongoing firing rate is smaller in AI state than TA state, it may be more energy efficient for the striatum to operate in the AI state. However, only in the TA state, weak stimuli can dramatically modulate the spectrum of correlations. The change in the bump structure implies that even a weak stimulus can introduce both positive and negative correlations. On a functional level such positive and negative correlations can initiate the spike timing dependent plasticity and form stimulus specific structure. Thus, we think that in TA state, not only the striatum can uniquely represent weak inputs but also can learn those inputs by changing synapses in an activity dependent manner.

### Model validation

Even though the three activity states of the network model capture various aspects in the striatal activity, it remains open question whether the striatal activity is organized in distinct NAs. *In vitro* experiments have provided some evidence for the NAs in the striatum [7]. The *in vivo* evidence is scarce and indirect at best. For example, the striatal activity in behaving animals is organized in spatially distributed NAs [6] and neighboring neurons are co-activated during the same task phase [4] indirectly points towards an assembly like organization of the striatal activity. But in most experiments done so far, neuronal activity was sampled too sparsely to make any clear statement about the spatial organization of the striatal activity. Recent advances in the optical recording methods allow for a measurement of activity of a large number of neurons located in a small neighborhood [37]. Such data can be used to validate the results of our network model. In addition, our model suggest that even the relationship between the firing rate and spike time irregularity (Fig. 2) can provide further indirect evidence for the existence of NAs.

The key to the emergence of transient NAs in the TA state is the non-monotonically shaped connection probability. Indirect estimates of the anatomical (from neuron morphologies) and functional connectivity both indicate that MSNs do not inhibit their nearest neighbors and the connection probability peaks at a distance of ≈ 80 *µ*m and then decays to zero beyond ≈ 200 *µ*m [19,20]. However, more experimental work is required to measure the spatial profile of not just the functional but also the structural connectivity within the striatum. However, more experimental work is required to measure spatial connection profile and spatially compact neural clusters that would support our network model. Alternatively, our models predicts that neurons participating in the NAs should have a low connection probability and share their inputs.

Recurrent networks with both excitatory and inhibitory (EI) neurons when interconnected according to a monotonically decaying (e.g. Gaussian) connectivity profile for excitation and inhibition, can be tuned to exhibit spatially clustered or stationary bump type activity [27]. A classical example are EI-networks in which the excitatory connectivity decays more rapidly with distance than the inhibitory connectivity. In such networks, the summation of excitatory and inhibitory connectivity kernels or excitatory and inhibitory synaptic strengths yield the well-known Mexican hat profile as the ‘effective’ connectivity kernel with its characteristic, non-monotonic shape. This EI network type shows the periodic activity bumps in which neighboring neurons become co-active because the local recurrent excitation is stronger than the local recurrent inhibition, whereas at larger distances, the recurrent inhibition exceeds the recurrent excitation. With such a connectivity profile, local recurrent excitation activates neighboring neurons which, in turn, inhibit the surrounding region because of the stronger distal inhibition. That is, in such EI-networks, a co-activated local group of neurons is brought together by their mutual, predominantly excitatory, connections and by its common field of surround inhibition. Examples of such behavior have been reported in the experimental literature, e.g. in the monkey prefrontal cortex [38]. By contrast, we found that for purely inhibitory networks, using both network simulations and neural field equations, a non-monotonic spatial connectivity kernel (such as the Gamma distribution) generates spatially clustered bump type activity for high background input. As shown by our mean field analysis, purely inhibitory networks with a Gaussian spatial connectivity profile cannot possibly show any spatially periodic bump activity. Intuitively, that is because co-activation of neighboring neurons requires that these neurons do not inhibit each other while creating an inhibitory surround. Thus, in purely inhibitory networks a co-activated local group of neurons is defined by the lack of mutual connectivity and the presence of a common range of surround inhibition. We would like to note that the stable bump state (WTA) observed in our networks closely resembles the grid patterns often observed in the medial entorhinal cortex of rodent and computational models of grid cells also use a non-monotonic kernel to form connections between inhibitory neurons [39,40]. In summary, here, we showed how the shape of the distance-dependent recurrent connectivity profile and the strength of ongoing, external excitation together determine the state of the ongoing activity as well as the stimulus response properties in a purely inhibitory network, such as the striatum. Thus, these results, when properly adapted to the specific inhibitory network of interest could provide important new insights into the functional characterization of the activity dynamics in inhibitory brain networks such as the striatum, the globus pallidus and the central amygdala.

## Models

### Neuron model

Neurons in the network were modeled as ‘leaky-integrate-and-fire’ (LIF) neurons. We intentionally chose this simple neuron model to ensure that the observed network dynamics would be the result of the connectivity profiles studied and not due to the intrinsic dynamics of a more complex neuron model. The subthreshold membrane potential dynamics of LIF neurons is given by:

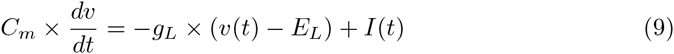

Synapses were modeled as conductance transients. The temporal profile of the transient conductance change in response to a single pre-synaptic spike was modeled as an alpha function. The recurrent connectivity within the inhibitory network such as the striatum [15–17] is weak and sparse. We adjusted the synaptic conductances to obtain weak synapses such that a unitary inhibitory postsynaptic potential (IPSP) had an amplitude of 0.8 mV at a holding potential of -44.0 mV and an excitatory postsynaptic potential (EPSP) had an amplitude of 1.6 mV at a holding potential of -70.0 mV. The various neuron and synapse parameters are summarized in Table 1. Whenever possible we used parameters corresponding to the striatal network [18].

**Table 1.**
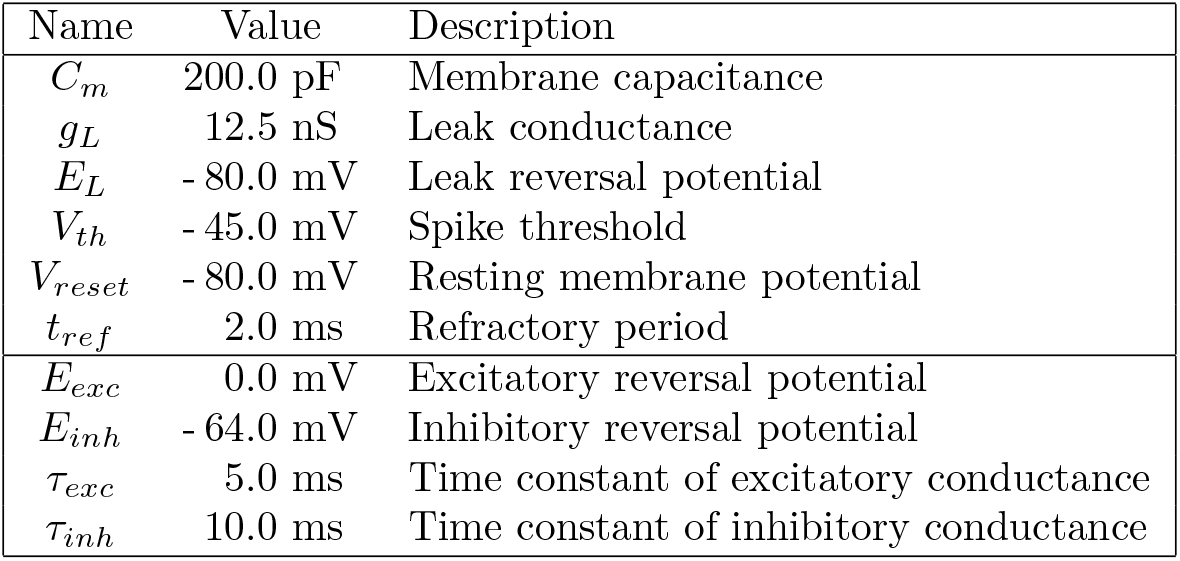
Parameter values

| Name | Value | Description |
| --- | --- | --- |
| $C_m$ | 200.0 pF | Membrane capacitance |
| $g_L$ | 12.5 nS | Leak conductance |
| $E_L$ | - 80.0 mV | Leak reversal potential |
| $V_{th}$ | - 45.0 mV | Spike threshold |
| $V_{reset}$ | - 80.0 mV | Resting membrane potential |
| $t_{ref}$ | 2.0 ms | Refractory period |
| $E_{exc}$ | 0.0 mV | Excitatory reversal potential |
| $E_{inh}$ | - 64.0 mV | Inhibitory reversal potential |
| $\tau_{exc}$ | 5.0 ms | Time constant of excitatory conductance |
| $\tau_{inh}$ | 10.0 ms | Time constant of inhibitory conductance |

Parameter values for the neuron and synapse model.

### Network architecture

We considered a population of 10,000 inhibitory neurons. The neurons were placed on a 100 × 100 grid and folded as a torus to avoid boundary effects. The distance between neighboring nodes in the grid network amounted to 10 *µ*m. Each neuron sent 1,000 inhibitory recurrent inputs to other neurons within the network (i.e. 10 % total connection probability), mimicking the sparse connectivity in most biological inhibitory networks. We implemented no self-connections within the network and neurons were allowed to form multiple connections.

To implement a distance-dependent connectivity we chose two qualitatively different spatial profiles [13]. For the first type of connectivity, we assumed that the distance-dependent connection probability decreased monotonically as a function of distance. To implement such connectivity we used the Gaussian distribution to model the distance-dependent decrease in connectivity (see Fig. 1a, top). We will refer to this network type as a *Gaussian network.*

For the second type of connectivity, we assumed that the distance-dependent connection probability decreased non-monotonically as a function of distance. Recent experimental data suggest that the connectivity between pairs of medium spiny neurons (MSNs) in the striatum may indeed change non-monotonically as a function of distance [19,20]. To implement such connectivity we used a Gamma distribution to model the distance dependent decrease in connectivity (Fig. 1a, bottom). We will refer to this network type as a *Gamma network.*

To appropriately compare these different network types we ensured that connection weights and the degree distributions were identical in both network types.

### Ongoing/Background excitatory drive

All neurons received background excitatory inputs mimicking the global ongoing activity in striatum that could either be due to cirtical or thalamic input or represent the alternation in striatal network dynamics due to an internal change (i.e. change in lateral inhibition). The external drive was modeled by uncorrelated homogeneous Poisson type spike trains with fixed firing rate. Each neuron received an independent realization of such spike train, obtained from the same underlying Poisson process. The rate of the Poisson type spike trains was varied systematically to study the different dynamical states of the inhibitory networks.

### Stimulus

To measure the impact of the stimulus response on network activity in each dynamic states we stimulated the network with an external stimulus, in addition to the background inputs described above. The stimulus provided to a subset of ≈ 45 neurons. To distribute the stimulated neurons in a spatial manner we defined a region of interest (ROI) of size 30 x 30 neurons. In the ROI we defined a location as centers for stimulated neurons chosen according to a two-dimensional Gaussian (*σ*: 2 grid points). The external stimulus was modeled as an injection of direct current to the selected neurons. The stimulus presentation lasted for one seconds and its amplitude was systematically varied between 0 – – 150 pA. To collect sufficient data for further statistical analysis each stimulus was presented 100 times to its assigned subset of neurons.

To estimate the flexibility of the ongoing activity dynamics for encoding stimuli we extended the simulation of the network with two different external stimuli. Each of the two external stimuli was provided to two non-overlapping subsets of ≈ 45 neurons. In the ROI, as described above, we defined two locations that were 10 grid points apart. Using these two locations as centers, stimulated neurons were chosen according to a two-dimensional Gaussian (*σ*: 2 grid points). The two sets of stimulated neurons are marked with red and orange clouds respectively in Fig. 5a). The two stimuli were presented alternatively (see Fig. 5a, bottom trace) and each stimulus presentation lasted for one second. Thus, the two stimuli differed only in terms of their target neurons, other properties (strength, duration) were identical.

### Characterization of the network dynamics

The network activity states were characterized on the basis of the spiking activity using standard descriptors: firing rates and the coefficient of variation of inter-spike intervals (*CV*_*ISI*_), the latter to measure the degree of irregularity of the spiking of individual neurons [21].

#### Entropy

To test whether the network activity was uniformly distributed over all neurons we measured the entropy of the network activity. The entropy was given by:

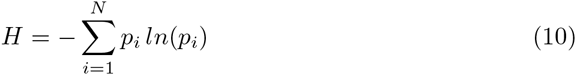

where *p*_*i*_ is the probability of neuron i to spike within a time window of 100 ms and *N* is the total number of neurons in the network. The lower the entropy, the more information is contained in the network activity.

#### Count and duration of activity bumps

We constructed a series of binary matrices representing the spatial map of the network activity over disjunct time windows of 100 ms each from the spike train recordings of all neurons. To enhance the spatial contrast, these matrices were convolved with the two-dimensional ‘Mexican hat’ shaped kernel (‘Ricker wavelet‘). The integral of the ‘Mexican hat’ kernel was set to zero to ensure that the filtered activity was comparable for different inputs. Additionally, the integrals of the positive and the negative part of the convolution kernel were set to 1 and -1, respectively. Subsequently, for each time step, the positions of the bump centroids were determined by detecting the local activity peaks. The positions of individual bumps were then used to compute the number of bumps and to determine the affiliation of neurons to individual bumps. Finally, trackers of time-merged bumps were used to analyze the life-span of the bumps, determined by finding similar positions of bump centroids over subsequent time frames.

### Characterization of the stimulus response

To estimate the influence of the external stimuli on striatal ongoing activity we analyzed the activity of 900 neurons in the ROI (of size 30 × 30 grid) surrounding the center of the stimulated region. Thus, we included both stimulated and unstimulated neurons in our analysis. To characterize the impact of external stimulus on striatal activity dynamics we measured the principal components (PC) of the activity and the spectrum of pair-wise correlations.

We analyzed the temporal evolution of the spike count using a sliding time window (window size: 100 ms; sliding step: 10 ms). Each 100 ms window resulted in a 900 dimensional array of spike counts. Thus, we reduced the 900 dimensional spiking activity sampled at a temporal resolution of 10 ms to a two-dimensional time-series of PCs. For each stimulus presentation (that lasted for 1 sec.) we measured the average of the first two PCs. These averaged PCs were plotted against each other and the scatter revealed how much the responses to the two stimuli differed from each other and from the ongoing activity.

The correlation spectrum was estimated for the activity of individual neurons. Neuronal activity was binned using a rectangular bin of 100 ms. We estimated the correlation spectrum for the trial-by-trial average of both ongoing and stimulus induced evoked activity. To estimate correlations, we considered 1000 ms long epoch of ongoing and evoked activity.

### Simulation Tools

All simulations of the networks of spiking neurons were performed using the NEST simulation software (http://www.nest-initiative.org) [22]. The dynamical equations were integrated at a fixed temporal resolution of 0.1 ms. Simulation data were analyzed with Python using the scientific libraries SciPy (http://www.scipy.org) and NumPy (http://www.numpy.org), and visualized using the plotting library Matplotlib (http://matplotlib.org).

## Supporting Information

**Figure 1. S1:**
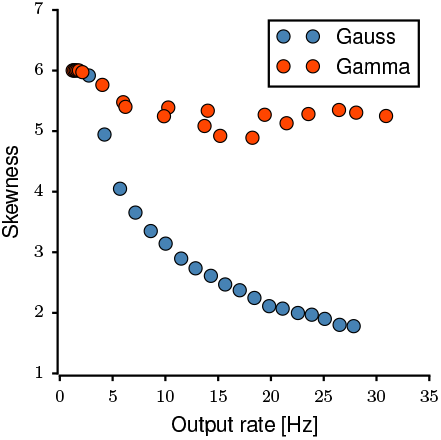
Skewness of firing rate distributions. The figure shows the skewness of the distribution of firing rates for different connectivity kernels as a function of the mean output rate. The skewness for the Gaussian kernel drops to the value 2 with increasing output rate, whereas the skewness for the Gamma kernel remains at a high value (> 5).

**Figure 2. S2:**
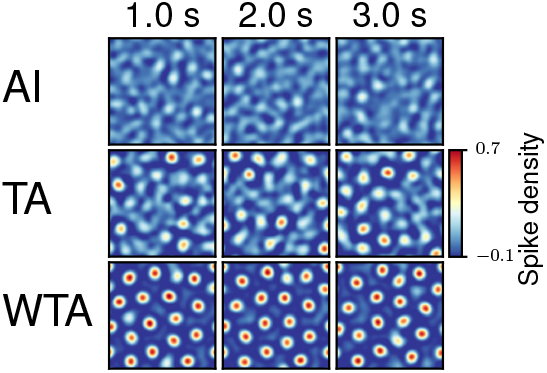
Characterization of the network dynamics for the bump activity state. Time series snapshots of the spatial pattern of bump activity, contrast-enhanced by Mexican hat filtering. A higher density of activated neurons forms a spatially localized activity bump. The time window for each frame is 100 ms. **Asynchronous irregular (AI)** No bump activity is observed the network activity remains noisy. **Transition activity (TA)** The network is in an unstable state, with several bumps appearing transiently, in the company of noisy activity. **Winner-takes-all (WTA)** The network forms mostly persistent bumps throughout the entire network.

**Figure 3. S3:**
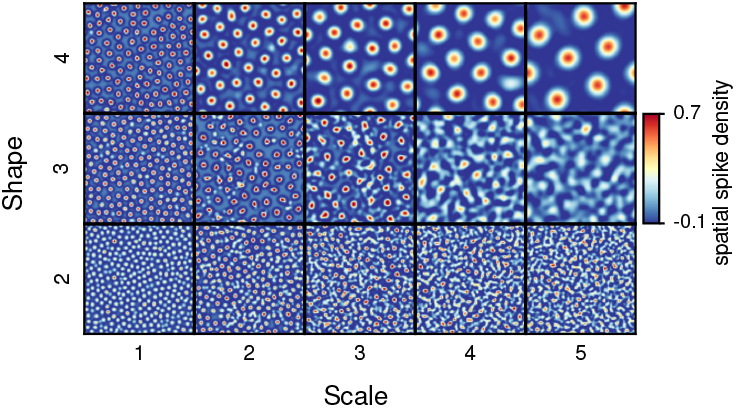
Spatial map of bump activity patterns for different Gamma distributions. A variety of Gamma values defines the size of bumps and the distance between bumps. This result illustrates the variety of bump capacity of the network, depending on the bump parameters. The rate of the external Poisson input here was 5 kHz.

**Figure 4. S4:**
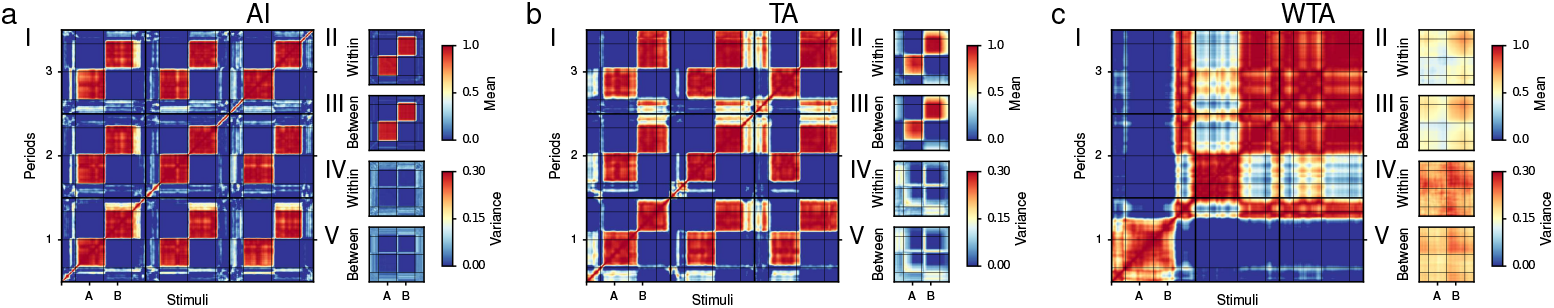
Temporal correlation of spiking activities in the AI state (a), TA state (b) and WTA state (c). The dynamic state of the network determines the impact of stimuli on the network activity. In each state, the bigger subplot (I) shows an example of temporal correlations for the first 3 out of 20 stimulus trials with a colormap that refers to a correlation score between 0 (blue) and 1 (red). The grid lines define switches of stimulus inputs (A,B,0). In each state, 4 smaller subplots (II-V) represent means and variances of temporal correlation for 20 trials. The first two subplots (II,III) show the mean of the temporal correlation within (diagonal) and between sequences which document the repeatability of the network response to the stimuli. The bottom two smaller subplots (IV,V) show the variance of the temporal correlation within (diagonal) and between sequences which document the variability of the network response to the stimuli.

## Acknowledgments

The authors acknowledge helpful discussions with Drs. Robert Schmidt, Ioannis Vlachos and Alejandro Fernandez-Bujan.

